# Three-dimensional shape cues affect human and artificial recognition systems differently

**DOI:** 10.64898/2025.12.01.691716

**Authors:** Mikayla Cutler, Luke Baumel, Joseph Tocco, William Friebel, George K. Thiruvathukal, Nichoals Baker

## Abstract

Humans and neural networks use shape and texture information differently. While shape is the primary cue in human object recognition, neural networks are more biased towards texture cues. Many tests of shape vs. texture bias have focused on shape recognition from an object’s external contour. However, shape information is also conveyed through internal contours, shading, and attached shadows, especially when an object is viewed from noncanonical perspectives. Using models from ShapeNet, we created datasets of 120,000 texture-substituted images of objects from many viewpoints with and without shading and attached shadows. We tested humans’ and several neural networks’ ability to classify these objects by both their shape and their texture. Humans were much better at classifying texture-substituted objects by their shape than any network, although these differences were greater when shape was defined only by the external contour than when 3D cues were included. Our findings suggest that networks’ texture bias is reduced when 3D cues are included in images. We next tested whether the inclusion of 3D cues benefitted humans and neural networks more for images of objects viewed from canonical or noncanonical perspectives. Consistent with earlier research, we found that 3D cues primarily benefitted humans for noncanonical images. For neural networks, the greatest performance gains were for canonical images. These findings suggest fundamental differences in how humans and networks use shading and attached shadows for object recognition. We argue that humans use these cues to infer objects’ 3D structures while neural networks use them as another surface-level cue like texture.

---

Deep neural networks (DNNs) match human performance on a variety of visual tasks (Attarian, 2020; Lou et al., 2022; Peterson et al., 2016; Prashnani et al., 2018; Sanders & Nosofsky, 2020; Yan et al., 2022; Zhang et al., 2018; Zhao et al., 2024) and predict neurophysiological activity in the visual brain (Adeli et al., 2023; Hong et al., 2016; Kar et al., 2019; Kar & DiCarlo, 2021; Storrs et al., 2021; Yamins et al., 2014). These successes have stirred interest in networks’ usability as image-computable models of visual perception. However, there are several key differences in the way humans and deep networks process visual information that have raised caution about their plausibility as models of visual cognition (Baker et al., 2023; Bowers et al., 2023; Doerig et al., 2020; Lonnqvist et al., 2025; Malhotra et al., 2023a; Younesi & Mohsenzadeh, 2024).

Among the most fundamental differences between humans and deep networks is in their use of shape. While humans are sensitive to the shape of objects above any other visual cue (Biederman & Ju, 1988; Elder & Velisavljević, 2009; Landau et al., 1988; Xu et al., 2004), networks are more reliant on the texture of the object for categorization (Azad et al., 2021; Baker et al., 2018; Geirhos et al., 2018; Heinert et al., 2024; Hermann et al., 2020; Islam et al., 2021; Iwata & Okuda, 2024; Jang & Tong, 2024). The inclusion of deceptive texture hurts DNN object recognition considerably more than human recognition (Baker et al., 2018; Geirhos et al., 2018; Hermann et al., 2020). Networks’ texture bias has been reduced by alternative training that limited the diagnostic value of texture information (Azad et al., 2021; Geirhos et al., 2018; Hermann et al., 2020; Jang & Tong, 2024; Tartaglini et al., 2022) or boosted the shape signal (Heinert et al., 2024). Whether these models leverage visual information like humans remains uncertain (Baker et al., 2020; Baker & Elder, 2022).

In studies where shape and texture information are put in direct competition with each other, a common approach is to remove the texture from an image of the original object and then substitute the texture from a different object onto the original object (Baker et al., 2018; Geirhos et al., 2018). Texture substitution at the image level can be done by converting color images of an object to grayscale and taking the elementwise product of the grayscale image and a texture image (Fig. 1a). Texture substitution can also be done by converting the object to a binary silhouette and substituting the figural region with a different texture (Fig. 1b). The former approach does not fully remove texture cues from the object’s shape, as can be seen from the preserved features in the eyes and face of the cat in Fig. 1a, which are the result of grayscale texture intensity (Burgert et al., 2025). The latter approach removes all texture but also any 3D shading information contained within the shape’s bounding contour.

**Figure 1.**
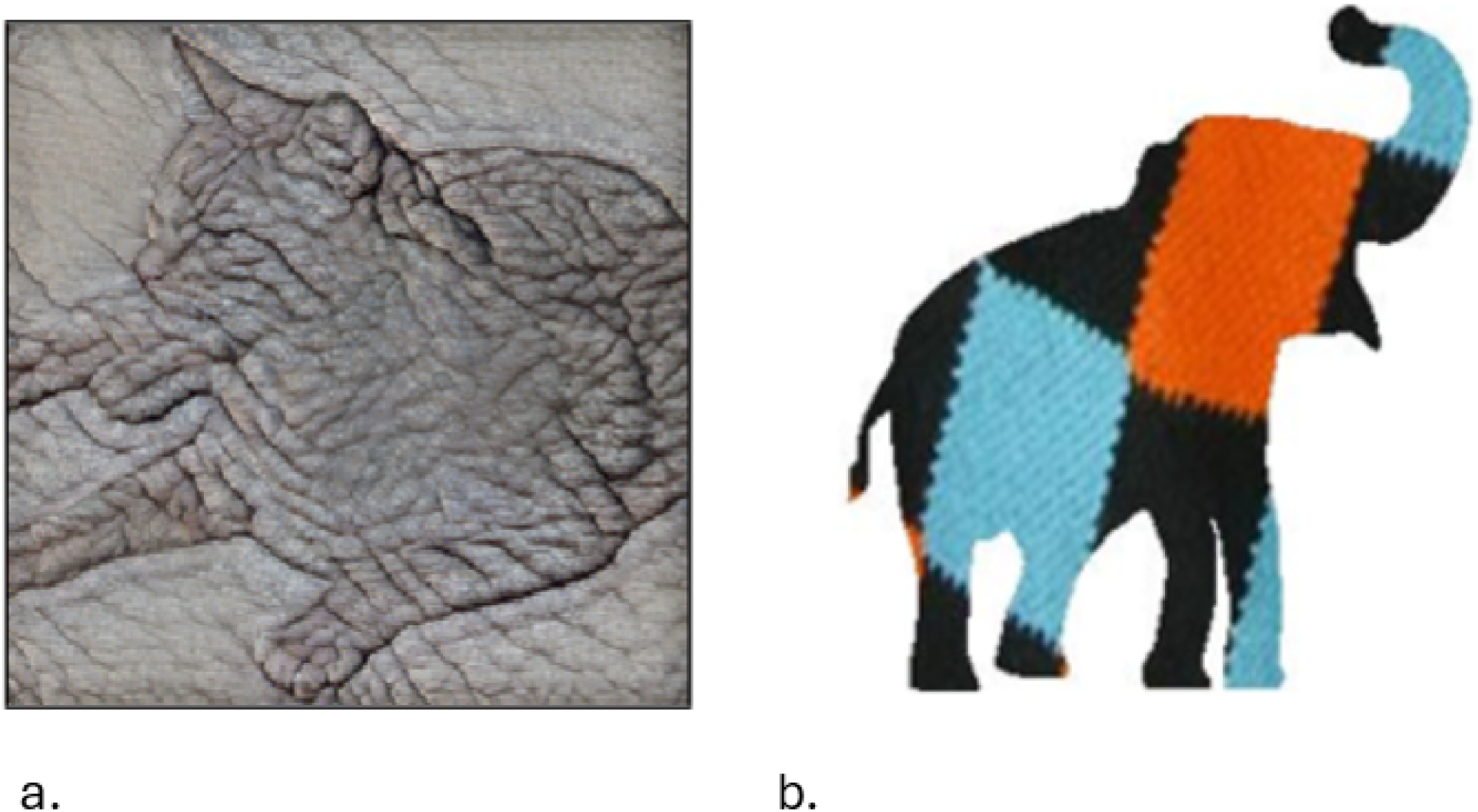
Texture substitution examples. a) Cat shape substituted with elephant texture. Reprinted with permission (Geirhos et al., 2018). b) Elephant shape substituted with argyle sock texture. Reprinted with permission (Baker et al., 2018).

Because of limitations in these approaches, it remains unstudied how shading and *attached shadows* (i.e., shadows cast from one part of an object onto another part of the same object) affect object recognition in DNNs. If networks recognize shapes more easily when shadows are preserved, one possibility is that their texture bias is caused not just by a preference for texture information but also by the loss of important shape 3D information conveyed by shading and attached shadows. Another possibility is that shading and attached shadows are treated as a texture cue by deep networks and are not leveraged by DNNs to infer objects’ 3D structures.

These two possibilities have also been considered for humans’ use of shading and attached shadows for object recognition. A great deal of information can be inferred about an object’s 3D structure based on shading and shadows (Cavanagh & Leclerc, 1989; Kriegman & Belhumeur, 1998), and humans may make use of these cues to form structural representations of objects (Bülthoff et al., 2023; Cavanagh & Leclerc, 1989; Favelle et al., 2017; Langer & Zucker, 1994). On the other hand, it has been argued that shadows are primarily used for recognition as an image-level cue, not for the inference of 3D structure (Cutzu & Edelman, 1994; Tarr et al., 1998).

We tested the interaction between shading and texture in both humans and DNNs by creating a novel dataset from 3D ShapeNet models overlaid with nondiagnostic texture. This allowed us to systematically manipulate texture information while preserving objects’ shape and attached shadows. We first compared humans’ and neural networks’ shape vs. texture bias with and without shading or attached shadows, which we refer to as *3D cues*. We predicted that humans, who are highly sensitive to the bounding contour of objects (Biederman & Ju, 1988; De Winter & Wagemans, 2004) would benefit only marginally from the inclusion of 3D cues, but that DNN performance would improve substantially with 3D cues included.

To the extent that shadows do benefit shape recognition in humans and deep networks, we also sought to understand why they are beneficial. Do shadows aid recognition because they increase the image-level similarity between a novel instance of an object and previously seen images, or do they allow for inference about an object’s 3D structure?

We tested this by comparing the performance advantage conferred by shadows for objects viewed from a typical (*canonical*) perspective vs. objects viewed from an atypical (*noncanonical*) perspective. In human perception, object recognition is easier for objects viewed from canonical perspectives (Hayward, 1998; Lawson, 1999; Newell & Findlay, 1997; Palmer et al., 1981). Differences in recognition performance between canonical and noncanonical images are greater when the images include only the object’s external contour than when internal contours (Lawson & Humphreys, 1999; Tian et al., 2016) or 3D cues such as shading and attached shadows are also present (Hayward, 1998; Newell & Findlay, 1999).

Likely, the interaction between canonicality and the inclusion of internal contours and/or shading and attached shadows has to do with the formation of structural, volumetric representations of object shape. If deep neural networks benefit from the inclusion of 3D cues equally for canonical and noncanonical images, that would be evidence that they do not form volumetric shape representations like humans do. It would also demonstrate that the asymmetrical benefit of 3D information for canonical and noncanonical images observed in humans is inconsistent with the view that humans recognize objects based on purely image-level similarities. DNNs constitute an ideal observer model for this kind of object recognition, so differences between humans and neural networks imply a different computational strategy for object recognition.

## Methods

### Models tested

We tested state-of-the-art three network architectures, ResNet-50, ViT-16, and SWIN. Below, we briefly describe key differences between the three models.

*ResNet-50*: ResNet-50 (He et al., 2016) is a very deep convolutional neural network. While it functions like other deep convolutional networks, it includes “skip connections”, which makes it possible for the network to include many layers while preserving the error signal in gradient descent. As one of the most famous convolutional networks, ResNet has been trained with several alternative curricula in efforts to increase its shape bias. We tested ResNet with three training methods.

1. ImageNet: The most standard curriculum for DNNs is ImageNet (Deng et al., 2009), a database of 1.2 million natural images of objects from 1,000 categories.
2. ImageNet and Stylized ImageNet: An alternative training method in which DNNs are trained on both standard ImageNet photographs and photographs from ImageNet that have undergone “style transfer”, which converts them into an artist’s painterly style. Doing so reduces the diagnostic value of texture and increases shape bias on some classification tasks (Geirhos et al., 2018).
3. Strong-blur: Another alternative training method aimed at simulating the visual experience of babies and vision in the periphery, two hypothesized causes of humans’ shape biases.

Blurred ImageNet images were convolved with a Gaussian blur kernel of varying sizes (σ = 0, 1, 2, 4 or 8 pixels, with equal probability). Shape bias was significantly greater in ResNet models trained with blurring than models trained only on the standard ImageNet dataset (Jang & Tong, 2024).

*ViT-16*: One of the first, and still among the most successful, transformer architectures applied to the object recognition task, ViT-16 (Dosovitskiy et al., 2021) uses a multi-head mechanism to learn the complex interactions between 16x16-pixel image patches for image classification. ViT greatly outperforms ResNet on shape classification tasks absent any diagnostic texture (Baker & Elder, 2022), possibly because its self-attention mechanism allows it to learn long-rage relations between parts of a shape.

Whereas convolutional networks are trained to learn local relations between nearby pixels in early layers, ViT is free to learn pixel relations between any patches in the image, regardless of distance. As a result, ViT needs much more training data than ReNet or SWIN (see below), but it performs very well when trained on large datasets. The model we tested was trained on JFT-300M, a large proprietary labelled image database, and finetuned on ImageNet for the 1,000 object categorization task.

*SWIN*: We also tested SWIN (Liu et al., 2021), a hierarchical transformer architecture that breaks images into small 4x4 pixel image patches in early layers but merges them into successively larger patches in deeper layers of the network. SWIN uses a local self-attention head which allows relations to be learned only between pairs of image patches in the same window of attention. This window of attention convolves over the whole image to learn different pixel relations. SWIN has more inductive biases than ViT and therefore requires less training data to accurately classify images. The model we tested was trained on ImageNet.

### Stimuli

#### 3D shape models

We selected 100 3D models from ShapeNet (Chang et al., 2015), a dataset of 3D shapes of both biological and non-biological objects. We chose ten object categories—five biological and five nonbiological—that were among the trained categories in ImageNet and had numerous high-quality models in ShapeNet. For each category, we selected 10 models. The biological categories were fish, elephant, butterfly, bird, and bear, and the non-biological categories were helmet, mailbox, bathtub, mug, and phone.

#### Texture images

For each object category, we also found 10 high-resolution texture images to pair with the shapes. Texture images were cropped so that they included no background pixels. *Testing datasets:* We created two datasets, one 3D and one 2D, upon which to test both humans and DNNs. We generated these datasets by first taking photographs of each of the 100 3D shape models twelve times in 30° increments for a total of 1,200 model images (100 shapes x 12 orientations). All models were photographed with white surfaces on a black background.

For the 3D stimuli, we created 100 retextured images for each model image by multiplying the grayscale pixel values of the image with the RGB values of each of the 100 texture images. We created stimuli for the 2D dataset by converting the same photographed images into binary images and multiplying each pixel value (one for the figure or zero for the background) with a texture image. This resulted in 120,000 3D and 2D retextured images that differed only in their inclusion of shading and attached shadow cues.

We hand-coded each object in our database as being viewed from a canonical perspective or a noncanonical perspective. Researchers have debated what determines whether a viewpoint results in a canonical or noncanonical image of an object. One view is that canonicality depends primarily on the visibility of features typically associated with a particular object, such as an elephant’s trunk or a piano’s keyboard (Palmer et al., 1981). Others have argued that canonicality depends only on the familiarity of a viewpoint (Blanz et al., 1996), although experiments where familiarity was controlled still showed canonicality effects (Edelman & Bülthoff, 1992). For our images, which are of familiar objects, these competing ideas make similar predictions. Objects viewed from a canonical perspective faced toward the viewer or in profile to the viewer. Objects viewed from a noncanonical perspective faced away from the viewer and often self-occluded distinctive features for recognition. An example canonical and noncanonical image from the 2D and 3D datasets is shown in Figure 2.

**Figure 2.**
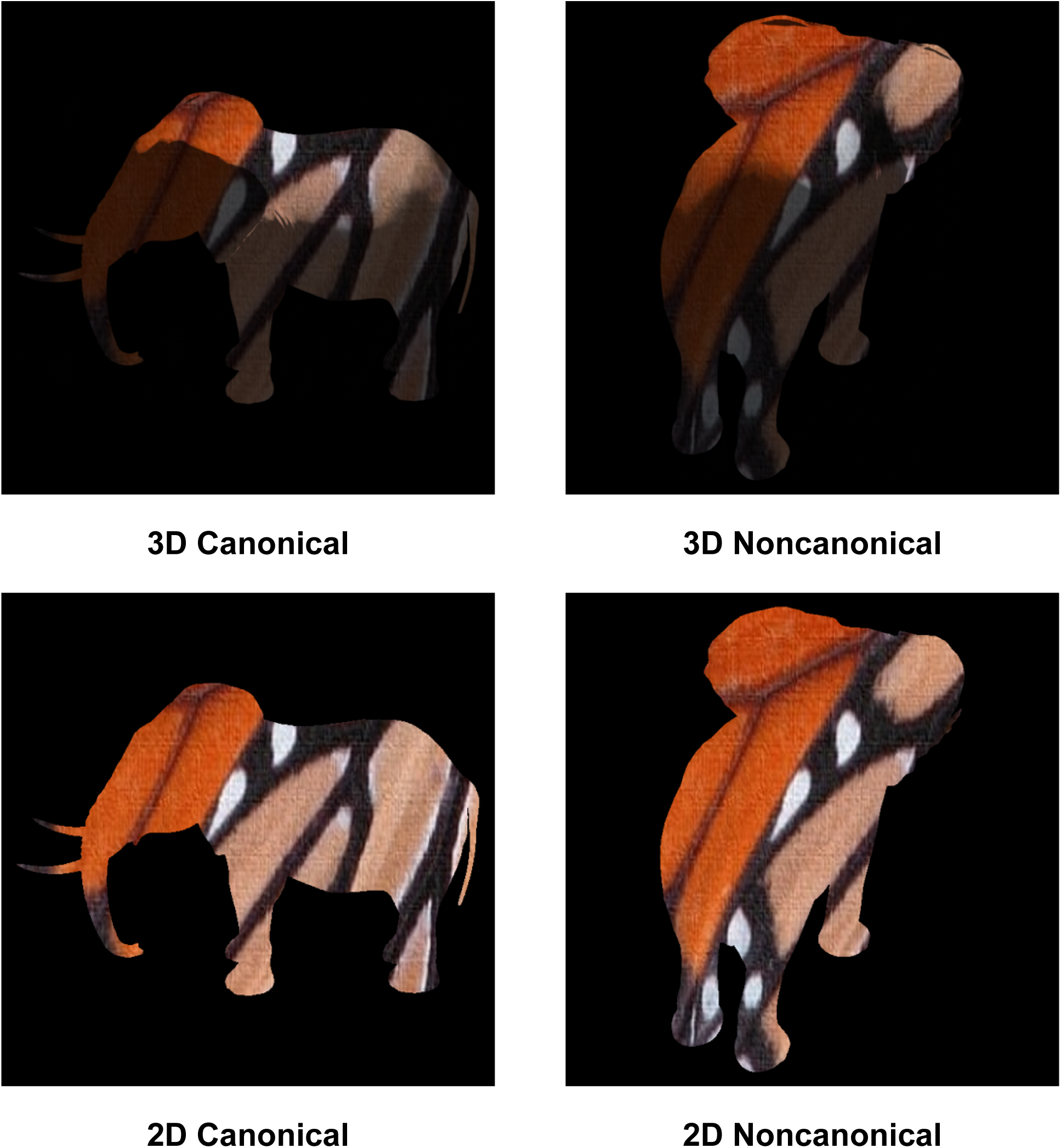
Sample images from Experiments 1 and 2. Top row: An elephant-butterfly with 3D information viewed from a canonical and noncanonical perspective. Bottom row: The same elephant-butterfly with only 2D information.

## Method

### Network experiments

We tested each DNN on each of the datasets described above. The 1,000 object categories networks are trained on with ImageNet are more granular than the entry-level categories we used to create our testing sets. We identified the subcategories in ImageNet that belonged to each entry-level category. The subcategories associated with each entry-level category are shown in Table 1. To determine networks’ classification decisions, we took the sum of all softmax values of subcategories that belonged to each entry-level category (Geirhos et al., 2018). The networks’ classification decisions were taken to be whichever entry-level category had the highest summed probability.

**Table 1.** Mapping of ImageNet categories to the ten entry-level categories used in our experiments.

|  |  |
| --- | --- |
| **Bathtub** | Bathtub, tub/vat |
| **Bear** | Koala, brown bear, black bear, polar bear, sloth bear, red panda, giant panda |
| **Bird** | House finch, snowbird, indigo finch, robin, bulbul, jay, magpie, chickadee, water ouzel |
| **Butterfly** | Ringlet butterfly, monarch butterfly, cabbage butterfly, sulphur butterfly, lycaenid butterfly |
| **Elephant** | Indian elephant, African elephant |
| **Fish** | Goldfish, crampfish, devilfish, eel, coho, rock beauty, anemone fish, sturgeon, garfish, lionfish, pufferfish |
| **Helmet** | Crash helmet, football helmet, gasmask |
| **Mailbox** | Mailbox/letter box |
| **Mug** | Coffee mug |
| **Phone** | Cellular telephone, dial telephone, payphone |

### Human experiments

We conducted two human experiments, one which compared humans’ sensitivity to shape and texture, and the other which compared humans’ ability to classify images’ shapes when object’s were viewed from canonical and noncanonical perspectives.

#### Participants

Forty-seven undergraduate students from Loyola University Chicago (32 women, 12 men, 3 nonbinary persons, 6 declined to report gender, *M*_age_ = 18.53) participated in both experiments for course credit. All participants had normal or corrected-to-normal vision and were naïve to the purpose of our experiments.

#### Experiment 1

Participants had two distinct tasks in Experiment 1. In the first 160 trials, they were instructed to identify an object’s shape from 10 categories, irrespective of its texture. We used 80 3D and the same 80 2D retextured images. Eight shapes from each category were randomly selected with a random texture and orientation with the caveat that the image’s texture and shape could never belong to the same category. Each object category was bound to a single key, and participants were instructed to press the key to identify the shape shown in the presented image. They were instructed to respond as quickly as possible without compromising accuracy. They completed 10 practice trials to familiarize themselves with the instructions and the mapping between object categories and response keys before beginning the main experiment.

Once they had completed the shape trials, participants were given a new set of instructions in which they were told to identify the texture belonging to the object in a presented image. As in the shape trials, stimuli were selected from the 2D and 3D retextured datasets. Eighty (eight textures for each of the 10 categories) images were selected from the dataset with shape randomly selected from the other nine categories and random orientation. Participants completed 10 practice trials before beginning the main experiment.

#### Experiment 2

In Experiment 2, participants’ only task was to identify the shape of the presented object, irrespective of texture. We had two independent variables, each with two levels.

The first factor we manipulated was shading cues. Images were either shaded by the three-dimensional structure of the object and had attached shadows (*3D*) or had no shading or attached shadows (*2D*). The second factor was canonicality of viewpoint. Images could be viewed from either a canonical or noncanonical perspective. Participants completed sixteen practice trials before beginning the main experiment. The main experiment consisted of 20 trials per condition.

### Dependent measures and analysis

#### Comparison of shape and texture sensitivity with and without 3D information

We compared networks’ texture bias with 2D vs. 3D images by comparing the proportion of trials in which the sum of subcategory probabilities was greatest for the object’s shape with proportion of trials in which the sum of probabilities was greatest for the object’s texture (see **Method**). For humans, we compared the proportion of correct trials when participants were tasked with identifying the object’s shape with the proportion of correct trials when they were tasked with identifying the object’s texture in Experiment 1. We also compared response time for shape identification trials with response time for texture identification trials. We tested for interactions between the identification task and presence of three-dimensional cues.

#### 2D vs. 3D object recognition from canonical and noncanonical perspectives

We hypothesized that if shading and attached shadows improve recognition by facilitating the formation of structural shape descriptions, then they should be of greater benefit to recognition of objects viewed from noncanonical perspectives than canonical perspectives. Systems that classify images based on image-level similarity would benefit from 3D information equally or more for images in which the object is seen from a canonical perspective.

## Results

### Shape and texture bias

The proportions of images that each DNN correctly classified by its shape and its texture are shown in Figure 3. We also calculated each network’s texture bias with and without 3D information included. Following Geirhos et al. (2018), we defined texture bias as the proportion of correct texture classifications divided by the proportion of correct shape or texture classifications. Values greater than 50% indicate a texture bias and values less than 50% indicate a shape bias.

**Figure 3.**
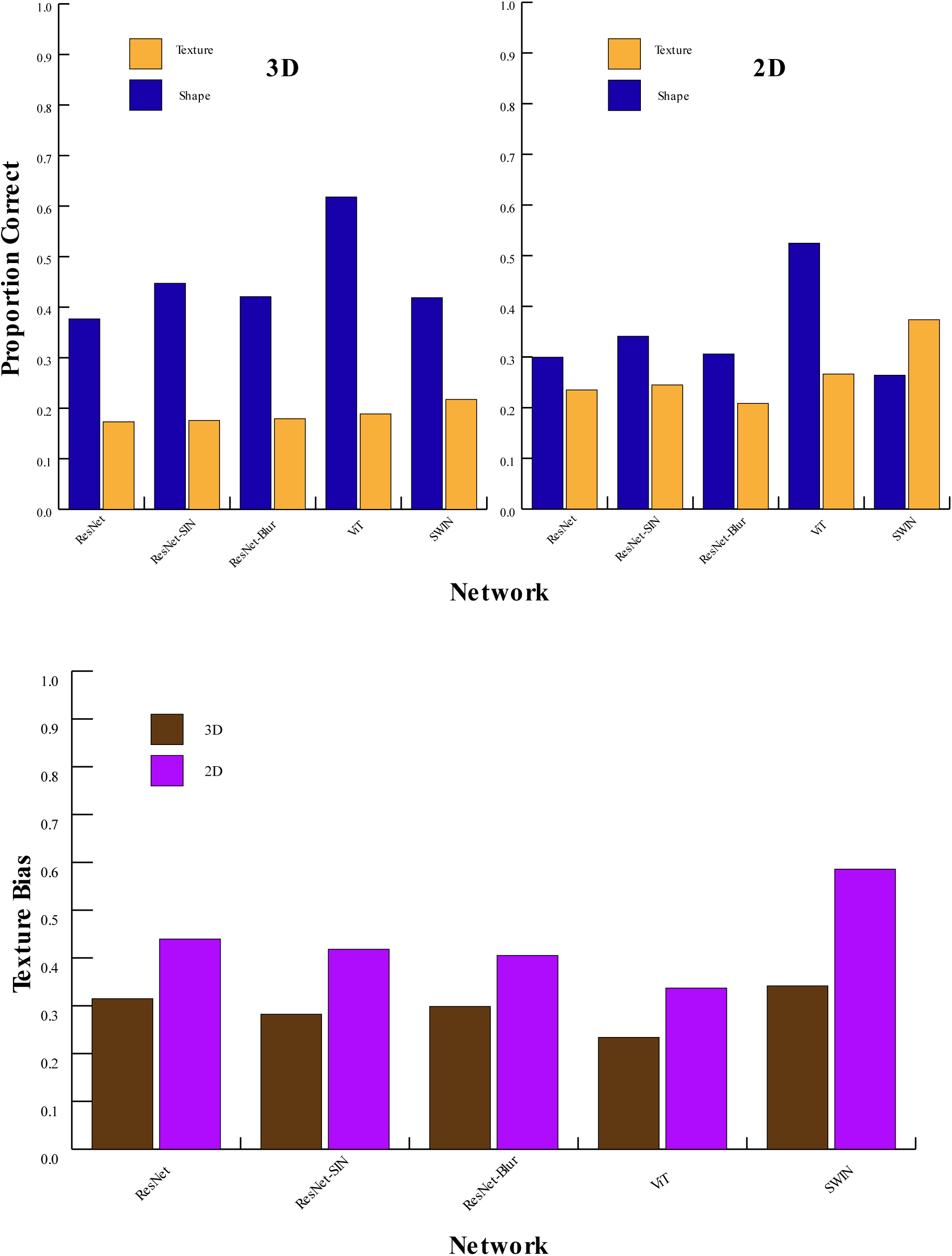
Neural network shape vs. texture results. Top row: Proportion of correct shape and texture classifications by each network with 3D and 2D information. Bottom row: Neural network texture bias in 3D and 2D, calculated as 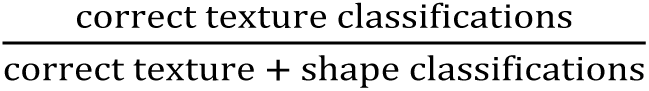

When 3D information was included, every network classified more images by their shape than by their texture. When shading and attached shadows were omitted (2D), this reversed for SWIN, and networks’ shape bias was reduced in all other networks. On average, networks’ shape classification accuracy was 11% better with shading and attached shadows included (46% vs. 35%) and their texture classification accuracy was 8% poorer with shading and attached shadows included (19% vs. 27%).

We also compared humans’ recognition of texture and shape. The results are shown in Figure 4. A 2x2 repeated measures ANOVA confirmed that humans responded significantly more accurately when tasked with classifying the object by its shape than by its texture (*F*(1,46) = 476.42, *p* < .001, η^2^_partial_= .91). They also responded more accurately when 3D information was included in shapes (*F*(1,46) = 16.21, *p* < .001, η^2^_partial_= .26). Paired comparisons found that classification accuracy was significantly better for shape than texture with (*t*(46) = 24.24, *p* < .001, *Cohen’s d* = 3.53) and without (*t*(46) = 16.13, *p* < .001, *Cohen’s d* = 2.35) 3D information.

**Figure 4.**
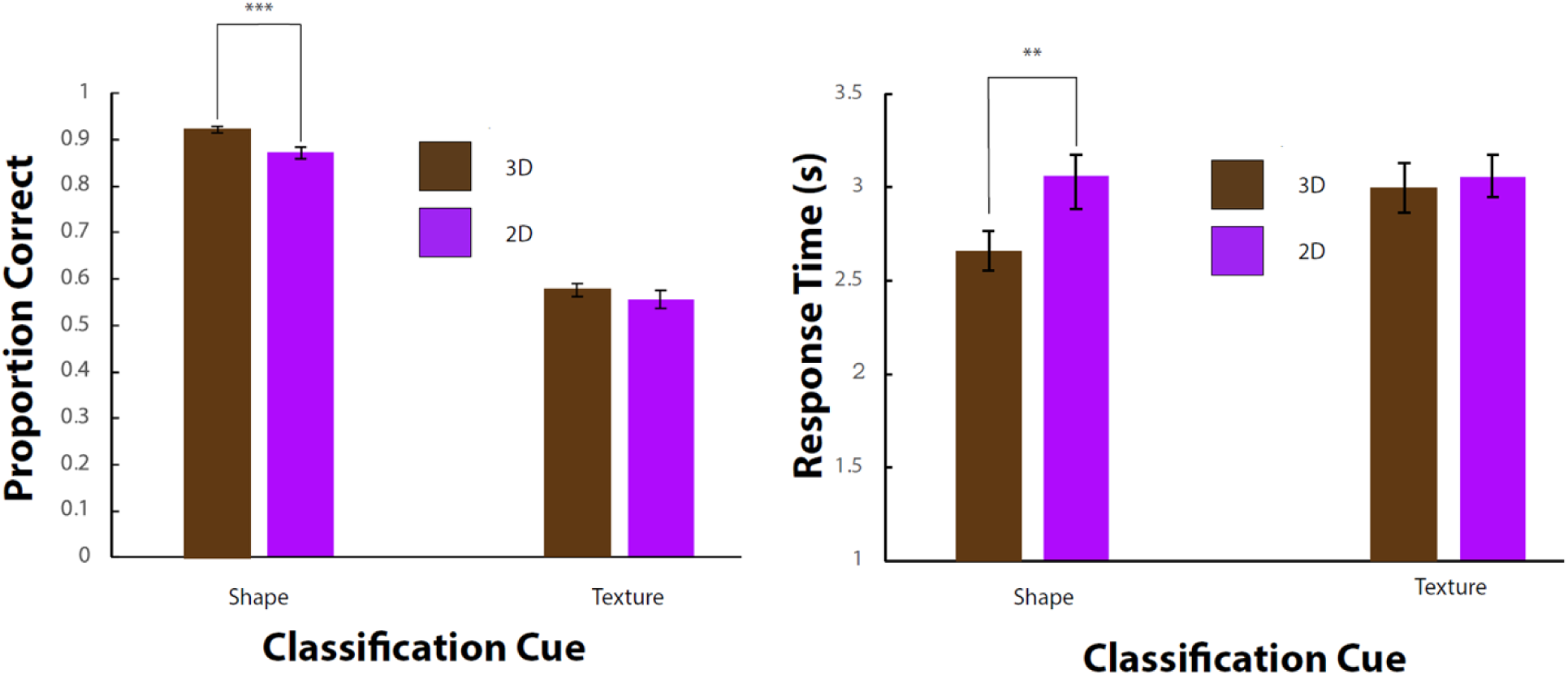
Human shape vs. texture results. Left: Classification accuracy for shape and texture classification tasks in 3D and 2D. Right: Mean response time for shape and texture classification tasks. Error bars reflect the standard errors of the mean.

We also found a significant interaction between classification cue and the inclusion of 3D information (*F*(1,46) = 4.60, *p* = .04, η^2^_partial_= .09). Participants recognized objects by their shape significantly more accurately with 3D information included (*t*(46) = 6.72, *p* < .001, *Cohen’s d* = 0.98) but did not significantly differ in their recognition of objects’ texture with or without 3D information (*t*(46) = 1.21, *p* = .23, *Cohen’s d* = .18). Response time results suggested no speed-accuracy tradeoff. Participants responded significantly more rapidly in shape trials with 3D information than in any of the other three conditions (all *t*’s > 2.60, *p*’s < .013, *Cohen’s d*’s > 0.38). Response times in other three conditions did not significantly differ (*t*’s < 0.74, *p*’s > .46, *Cohen’s d*’s < 0.11).

### Recognition of objects from canonical and noncanonical perspectives

We compared neural networks’ shape classification accuracy for objects viewed from a canonical perspective and a noncanonical perspective with and without 3D information in the image. The results are displayed in Figure 5. As reported above, shape classification performance improved with 3D information included. This improvement was no greater for images viewed from a noncanonical perspective than images viewed from a canonical perspective. In fact, all five networks improved more when 3D information was included in images taken from a canonical viewpoint than when it was included in images taken from a noncanonical viewpoint.

**Figure 5.**
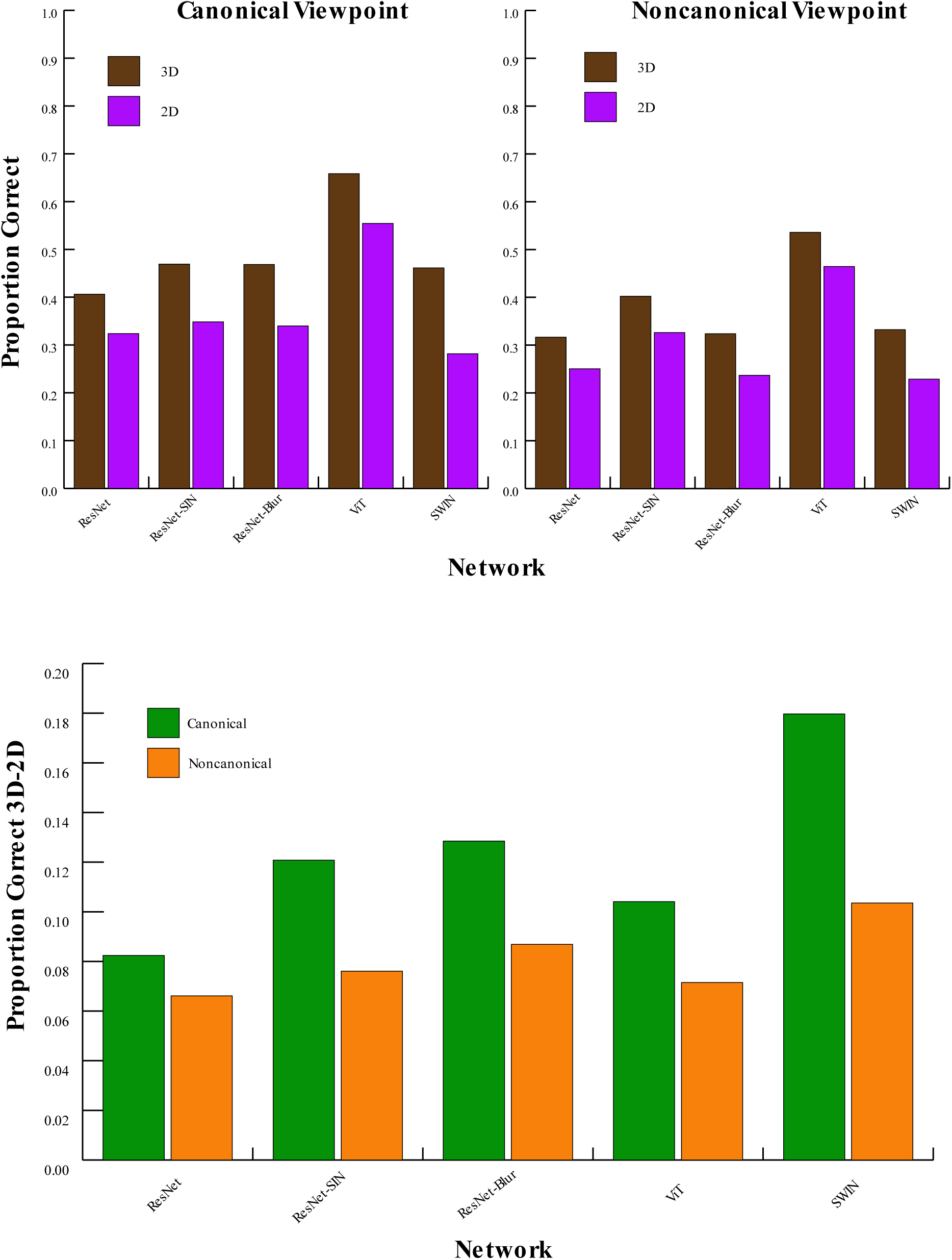
Neural network canonical vs. noncanonical shape recognition results. Top row: Network classification accuracy for images with 3D or 2D information viewed from a canonical or noncanonical perspective. Bottom row: Performance gain from the addition of 3D information in canonical and noncanonical images.

We also analyzed the human data from Experiment 2 where viewpoint and 3D information were manipulated. The results are shown in Figure 6. A 2x2 repeated measures ANOVA confirmed significant main effects for viewpoint canonicality (*F*(1,46) = 58.3, *p* < .001, η^2^_partial_= .56) and for the inclusion of 3D vs. 2D information (*F*(1,46) = 61.1, *p* < .001, η^2^_partial_ = 57).

**Figure 6.**
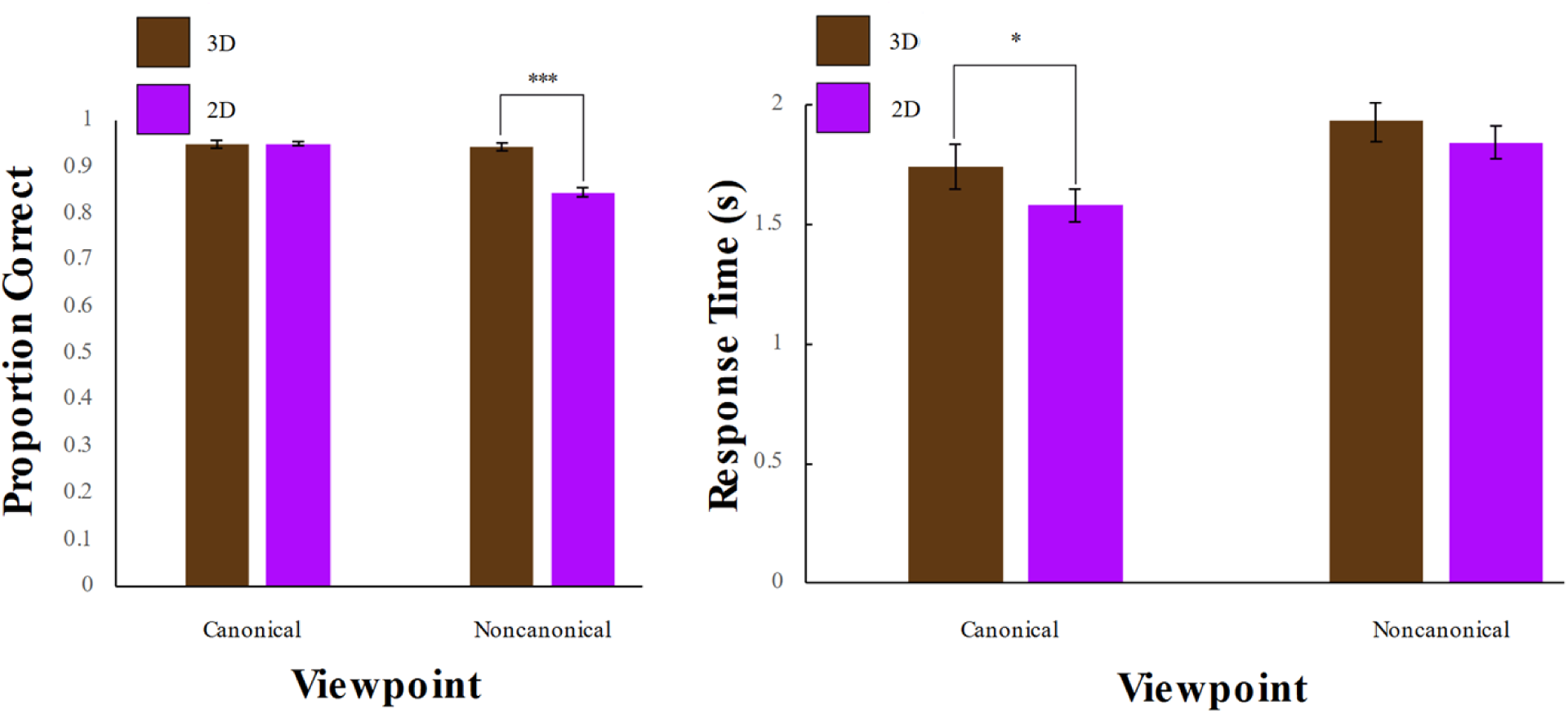
Human canonical vs. noncanonical shape recognition results. Left: Mean accuracy for images viewed from canonical vs. noncanonical perspectives with and without 3D information. Right: Mean response time. Error bars reflect the standard error of the means.

We also found a significant interaction between viewpoint canonicality and the inclusion of 3D information (*F*(1,46) = 59.9, *p* < .001, η^2^ = .57). The inclusion of 3D shape information did not significantly affect humans’ response accuracy for images viewed from a canonical perspective (*t*(46) = 0.21, *p* = .83, *Cohen’s d* = 0.03), but it did significantly improve human performance for images viewed from a noncanonical perspective (*t*(46) = 9.78, *p* < .001, *Cohen’s d* = 1.43). There is no evidence of a speed-accuracy tradeoff from the response time data. Humans’ equally accurate response rate for 2D images viewed from a canonical perspective was not a result of longer view time. In fact, humans responded significantly more rapidly in these trials (*t*(46) = 2.49, *p* = .02, *Cohen’s d* = 0.36). Differences in response time with 3D and 2D information were not significant for noncanonical images (*t*(46) = 1.73, *p* = .09, *Cohen’s d* = 0.25).

## Discussion

Deep networks are known to be more texture-biased than humans (Azad et al., 2021; Baker et al., 2018; Geirhos et al., 2018; Heinert et al., 2024; Hermann et al., 2020; Islam et al., 2021; Iwata & Okuda, 2024; Jang & Tong, 2024). However, previous work has not systematically manipulated the inclusion of 3D shape cues such as shading and attached shadows to test their effect on object recognition and their interaction with texture cues. We designed a novel testing set using 3D models from ShapeNet to measure the effect of including 3D cues on networks’ shape bias.

Consistent with previous work, we found that absent 3D cues from the object’s shape, neural networks are considerably more texture-biased than humans. In five top-performing networks, DNNs had an average texture-bias of 44% with only the external contour present. Networks classified objects correctly by their shape in an average of 35% of trials and by texture in 27% of trials. Humans classified objects correctly by shape in 87% of trials and by texture in 56% of trials without 3D cues.

The tests we used for neural networks and humans were not identical: In the neural network test, we presented a cue-conflict image and measured the probability the network assigned to both the shape and texture cue. In the human test, we explicitly instructed participants to classify images by shape or texture. These task differences cannot account for humans’ superior shape recognition performance with cue-conflict stimuli. Humans’ shape classification accuracy (87%) exceeded neural networks’ classification accuracy for both shape and texture (61%). This held true across all networks, so even if all the classifications that neural networks made for the object’s texture were counted as a correct shape classification, no DNN would have performed as well as humans on the cue-conflict stimuli.

While DNNs’ texture bias is greater than humans’, most networks were biased more towards shape cues than texture cues. All networks but SWIN classified more objects correctly by their shape than by their texture, even when 3D cues were omitted from the cue-conflict images. These findings challenge the view that contemporary DNNs rely primarily on texture for object recognition (Baker et al., 2018; Geirhos et al., 2019). While DNNs are much more influenced by texture than humans, shape still plays an at least equal role in classification decisions.

Comparing between DNNs with different architectures or training curricula, we found surprisingly small differences in texture-bias for ResNet trained only on ImageNet and ResNet trained to specifically reduce texture-bias, such as by augmenting ImageNet with stylized (Geirhos et al., 2018) or blurred (Jang & Tong, 2024) images. With only the external contour, ResNet trained on ImageNet had a texture bias of 44%, which went down to 42% and 41% for ResNet-SIN and ResNet-Blur, respectively. With 3D shape cues included, ResNet had a texture-bias of 31%, which went down to 28% and 30% in ResNet-SIN and ResNet-Blur. These differences may be meaningful, but they are relatively modest.

ViT was the least texture-biased network among the five we tested. Intriguingly, SWIN, the other tested transformer network, was the most texture-biased. One way ViT differs from SWIN is that it has a global self-attention mechanism that allows it to learn long-range relations between pixels. This could help the network learn diagnostic features that go beyond texture, which is locally defined. SWIN, whose self-attention mechanism is constrained to learn pixel relations within a local window, may be driven to learn more local cues. Another difference between ViT and SWIN is in the quantity of data upon which each network is trained. ViT was exposed to around 300 times more images than SWIN during training. Size of training data is a major factor in other global shape cues like contour integration (Lonnqvist et al., 2025), so it could be that the differences in ViT and SWIN’s training curricula, not their architectures, precipitated differences in their texture-biases.

All networks’ shape bias increased substantially when 3D cues like shading and attached shadows were included from the shape model. On average, DNNs’ shape bias increased from 56% to 70% with 3D cues included. Their shape recognition accuracy increased from 35% to 45%. Stimuli that put shape and texture in conflict with each other have not typically included 3D shape information because it is difficult to render separately from texture when starting from a photograph. By rendering 3D models, we preserved shapes’ shading and attached shadows while substituting all other surface properties with texture from another object.

Humans also benefitted from the inclusion of 3D cues, although not by as much. Shape recognition performance increased from 87% to 92% with the addition of 3D cues. Part of humans’ smaller performance gain is likely due to their excellent ability to recognize shapes with only the bounding contour (Wagemans et al., 2008; Baker & Elder, 2022; Lloyd-Jones & Luckhurst, 2002).

We next compared the way in which 3D cues benefit recognition in biological and artificial systems. One reason that 3D cues might be beneficial is that they operate as another textural or image-level cue that is consistent with the object’s shape. Images in the testing set were more similar to images that networks were trained on, or that humans had previously seen, when they included 3D cues. Luminance differences caused by shading and attached shadows might function no differently than luminance differences caused by patterns of fur or differently colored feathers. Another possibility is that humans and/or deep networks make use of shading and attached shadows to infer a shape’s 3D structure. Under this hypothesis, shadows would not be beneficial to recognition because they increase the image-level similarity between a test image and a training image, but because they increase the probability that the structural representation formed from the test image matches representations formed from previous visual experiences.

In human perception, shading, shadows, and internal contours help most with the recognition of objects viewed from a noncanonical perspective (Hayward, 1998; Lawson, 1999; Newell & Findlay, 1997; Tian et al., 2016). For these images, there may not be enough information in the external contour to form a structural description of an object’s 3D shape (Palmer et al., 1981). Shading and attached shadows furnish additional information that aids with the formation of a 3D shape representation (Bülthoff et al., 2023; Favelle et al., 2017; Marr, 2010).

The human data in Experiment 2 replicated these previous findings. Humans performed similarly with and without 3D shape cues when objects were viewed from a canonical perspective (95% vs. 95%--no significant difference), but when objects were viewed from a noncanonical perspective, performance was significantly worse for images that did not include 3D cues than images that did (85% vs. 94%).

The same was not true in deep networks. On average, DNNs’ accuracy was 12% higher when 3D information was included in images viewed from a canonical perspective and only 8% higher when it was included in images viewed from a noncanonical perspective. For deep networks, shading and attached shadows seem to be beneficial for recognition only because they serve as another image-level cue for recognition. Objects viewed from a canonical perspective likely benefit from these 3D cues more because there are more images including those cues from that perspective in the training data.

These findings are consistent with a growing literature showing differences between humans’ and DNNs’ use of shape for object recognition. As the current work and other studies show, DNNs can classify objects based only on shape information (Baker et al., 2018; Baker & Elder, 2022; Kubilius et al., 2016). However, research suggests that shape-based classification in neural networks relies on local shape features, not more configural aspects of shape (Baker et al., 2018, 2020; Baker & Elder, 2022; Brendel & Bethge, 2019; Burgert et al., 2025; Malhotra et al., 2023b). The formation of a structural, volumetric representation of object shape and subsequent use of this representation for recognition would likely require a level of abstract and configural processing beyond the capabilities of current DNN models.

An alternative hypothesis is that neither humans nor neural networks form structural representations of an object’s shape. Possibly, humans benefit more from shading and attached shadows in noncanonical images because the object’s external contour is more ambiguous in these images and image-level shading information is more discriminative. The pattern of results from all five tested neural networks suggests this is not the case. DNNs serve as an ideal observer model for recognition based on image-level similarities. If shading and attached shadows were more discriminative for noncanonical images, then we should have found the same interaction between canonicality and the addition of 3D cues in neural networks that we observed in human observers. Instead, we found that DNNs benefitted more from the inclusion of 3D information in canonical images than noncanonical images, suggesting that shading is not simply more informative for noncanonically viewed objects at the image-level.

## Conclusion

Like many previous studies, the current work found that neural networks are substantially more texture-biased than humans. However, this texture bias is reduced when shape cues like shading and attached shadows are included in an image along with the object’s external contour. While both humans and DNNs benefit from the inclusion of 3D shape cues, they are not alike in the way these cues benefit object recognition. Human performance primarily improves for images viewed from a noncanonical perspective where the external contour may be insufficient to accurately represent the object’s 3D structure, while DNNs benefit most for images of objects from familiar viewpoints where they may have had more exposure to image-level luminance patterns. These results suggest that shading and attached shadows help humans form structural, three-dimensional representations of an object while they help DNNs only with classifying objects according to image-level similarities.

